# Structure and Functional Binding Epitope of V-domain Ig Suppressor of T-cell Activation (VISTA)

**DOI:** 10.1101/597716

**Authors:** Nishant Mehta, Sainiteesh Maddineni, Irimpan I. Mathews, Andres Parra Sperberg, Po-Ssu Huang, Jennifer R. Cochran

## Abstract

V-domain Ig Suppressor of T cell Activation (VISTA) is an immune checkpoint protein that inhibits the T - cell response against cancer. Similar to PD-1 and CTLA-4, antibodies that block VISTA signaling can release the brakes of the immune system and promote tumor clearance. VISTA has an Ig-like fold, but little is known about its structure and mechanism of action. Here, we report a 1.85 Å crystal structure of the human VISTA extracellular domain and highlight structural features that make VISTA unique among B7 family members. Through fine-epitope mapping, we also identify solvent-exposed residues that underlie binding to a clinically relevant anti-VISTA antibody. This antibody-binding region is also shown to interact with V-set and Ig domain-containing 3 (VSIG3), the recently proposed functional binding partner of VISTA. The structure and functional epitope determined here will help guide future drug development efforts against this important checkpoint target.

## Introduction

V-domain Ig Suppressor of T-cell Activation (VISTA) is an immune checkpoint protein involved in the regulation of T cell activity. VISTA is highly expressed on myeloid-derived cells such as CD11b+ monocytes, CD11c+ dendritic cells, and to a lesser extent on CD4+ and CD8+ T cells^1^. Similar to the well-studied PD-1, PD-L1, and CTLA-4 checkpoint proteins, the presence of VISTA results in reduced T cell activation and proliferation. The mechanism of action for this effect, however, is unclear as VISTA has been thought to function as both a ligand and a receptor. As a ligand, VISTA is expressed on antigen-presenting cells and binds an unknown receptor on T cells to inhibit downstream T cell activation^1,2^. As a receptor, VISTA is expressed on T cells and transduces intracellular inhibitory signals after ligand binding to curtail T cell activity^3,4^. A proposed ligand for VISTA has recently been identified as V-Set and Immunoglobulin domain containing 3 (VSIG3)^5^.

Checkpoint proteins have been found to be overexpressed by cancer cells or their surrounding immune cells and prevent anti-tumor activity by co-opting natural regulation mechanisms to escape immune clearance. In particular, VISTA is upregulated on tumor infiltrating leukocytes, including high expression on myeloid-derived suppressor cells (MDSCs)^6,7^. Through VISTA signaling, these inhibitory immune cells prevent effective antigen presentation and indirectly promote tumor growth. VISTA is implicated in a number of human cancers including skin (melanoma)^8^, prostate^9^, colon^10^, pancreatic^11^, ovarian^12^, endometrial^12^, and lung (NSCLC)^13^. Additionally, VISTA levels have been found to increase after anti-CTLA-4 treatment (ipilimumab) in prostate cancer^9^ and after anti-PD-1 treatment in metastatic melanoma^8^, highlighting VISTA expression as a method of acquired resistance to currently available checkpoint inhibitors. For these reasons, VISTA is an important cancer immunotherapy target for drug development efforts.

The human VISTA protein is 279 amino acids in length, comprising a 162 amino acid extracellular domain, a 21 amino acid transmembrane domain, and a 96 amino acid cytoplasmic domain. The cytoplasmic domain lacks any immunoreceptor tyrosine-based signaling motifs, but does contain multiple casein kinase 2 and phosphokinase C phosphorylation sites that could play a role in signal transduction. Protein sequence analysis has clustered VISTA with the B7 family group of ligands (CD80, CD86, PD-L1, PD-L2, ICOSL, and CD276), all of which contain a conserved IgV-like fold. Among these proteins, VISTA is an outlier with relatively low sequence homology to other family members. The closest homolog within the B7 family is PD-L1, which shares only 22% sequence identity with VISTA. The VISTA extracellular domain contains two canonical cysteines that are conserved in Ig-like proteins, and also has four unique cysteine residues that are not present in other B7 family members. The low sequence homology and additional cysteine residues have hindered accurate structural modeling of VISTA based on sequence alone and present a clear need for a high resolution crystal structure.

Antibodies against VISTA have shown anti-tumor efficacy in multiple syngeneic mouse models^1,6,14^. Therapeutic development has progressed to human clinical trials with the development of anti-human VISTA antibodies led by Janssen Therapeutics. A purported lead anti-VISTA antibody (called ‘VSTB’ here) inhibits VISTA signaling *in vitro* and shows tumor regression in a MB49 syngeneic mouse model of bladder cancer^15^; however, little is known about its mechanism of inhibition. Putative regions of interaction between VSTB and VISTA have been proposed, but a specific binding epitope has not been identified. It is also unknown if the anti-VISTA activity of VSTB is derived from the blockade of VISTA/VSIG3 interaction. Moreover, although murine and human VISTA share 70% sequence homology, VSTB is not cross-reactive between species, which introduces challenges with testing in murine tumor models.

We present a high resolution crystal structure of the human VISTA protein and highlight distinct characteristics of the unique IgV-like fold that sets VISTA apart from other B7 family proteins. We also use combinatorial methods to map the VSTB/VISTA binding epitope, and further examine this region for potential VSIG3 interactions. Structural comparisons and epitope analyses performed here provide a blueprint for further VISTA mechanistic research and the development of next generation anti-VISTA therapeutics.

## Results

### Crystallization of the extracellular domain of human VISTA

The extracellular domain (ECD) of human VISTA, containing a C-terminal hexahistidine tag, was recombinantly expressed in human embryonic kidney cells and purified from the supernatant using immobilized metal affinity chromatography. The VISTA ECD was found to be hyper-glycosylated, producing a diffuse protein band that appeared to be ~15 kDa larger than its predicted molecular mass upon analysis by gel electrophoresis (SDS-PAGE). To generate well-formed crystals, we attempted to minimize glycosylation by introducing mutations to known N-linked glycosylation sites and also through enzymatic cleavage of sugars using a glycosidase. Analysis of the VISTA sequence highlighted five potential locations for N-glycan modification via a NXT/S motif. We mutated three of these asparagine residues (N59, N76, and N158) to glutamine. These selected positions were found to be three or more amino acid residues away from predicted secondary structure and therefore changes at these positions were less likely to affect native folding of the protein. In order to further reduce N-linked glycosylation, we added Kifunensine, a mannosidase I inhibitor, to the mammalian cell culture media prior to plasmid transfection. In addition, the purified VISTA protein was treated with the glycosidase Endo Hf prior to crystallization trials. These efforts resulted in improved discreteness and decreased apparent mass of the purified protein as compared to wild-type VISTA, indicating decreased glycosylation (Supp. Fig. 1).

Crystal trays were established in sitting drop format and placed at 12° C overnight. An optimized condition using seeds from prior, smaller crystal hits produced diffraction quality crystals. A complete dataset to 1.85 Å was collected by x-ray diffraction at the Stanford Synchrotron Radiation Lightsource (SSRL). VISTA does not have a suitable template for molecular replacement as no VISTA homologs are deposited in the PDB and the closest templates have sequence identity under 25%. The crystal structure was therefore solved using a combination of molecular replacement (MR), Rosetta modeling, and native sulfur single-wavelength anomalous diffraction (SAD) methods. Briefly, an iterative MR-Rosetta pipeline was used to find MR solutions, which were further rebuilt and refined with Rosetta. The model from the automated Rosetta procedures was then manually refined with Phenix to obtain the final structure.

The VISTA ECD contains three disulfide bonds comprising all six cysteine residues found in the VISTA sequence (Fig. 1A). The structure consists of ten beta strands and three alpha helices arranged in a canonical beta-sandwich formation (Fig. 1B). The protein is divided into two faces: six beta strands forming one coplanar surface and four beta strands comprising the other. The protein fragment between strands C and C’ is comprised of 21 residues forming an unstructured loop and four residues in a predicted alpha helix. Of the 25 residues in this region, six are predicted to be positively charged while only three are negatively charged, creating a net positive charge on this face of the protein. This positive plane is reflected in blue using the APBS electrostatic prediction tool (Fig. 1C). The structure presented here represents a common Ig-like fold, but as described below, closer examination reveals important differences that make VISTA unique among B7 family proteins.

**Figure 1.**
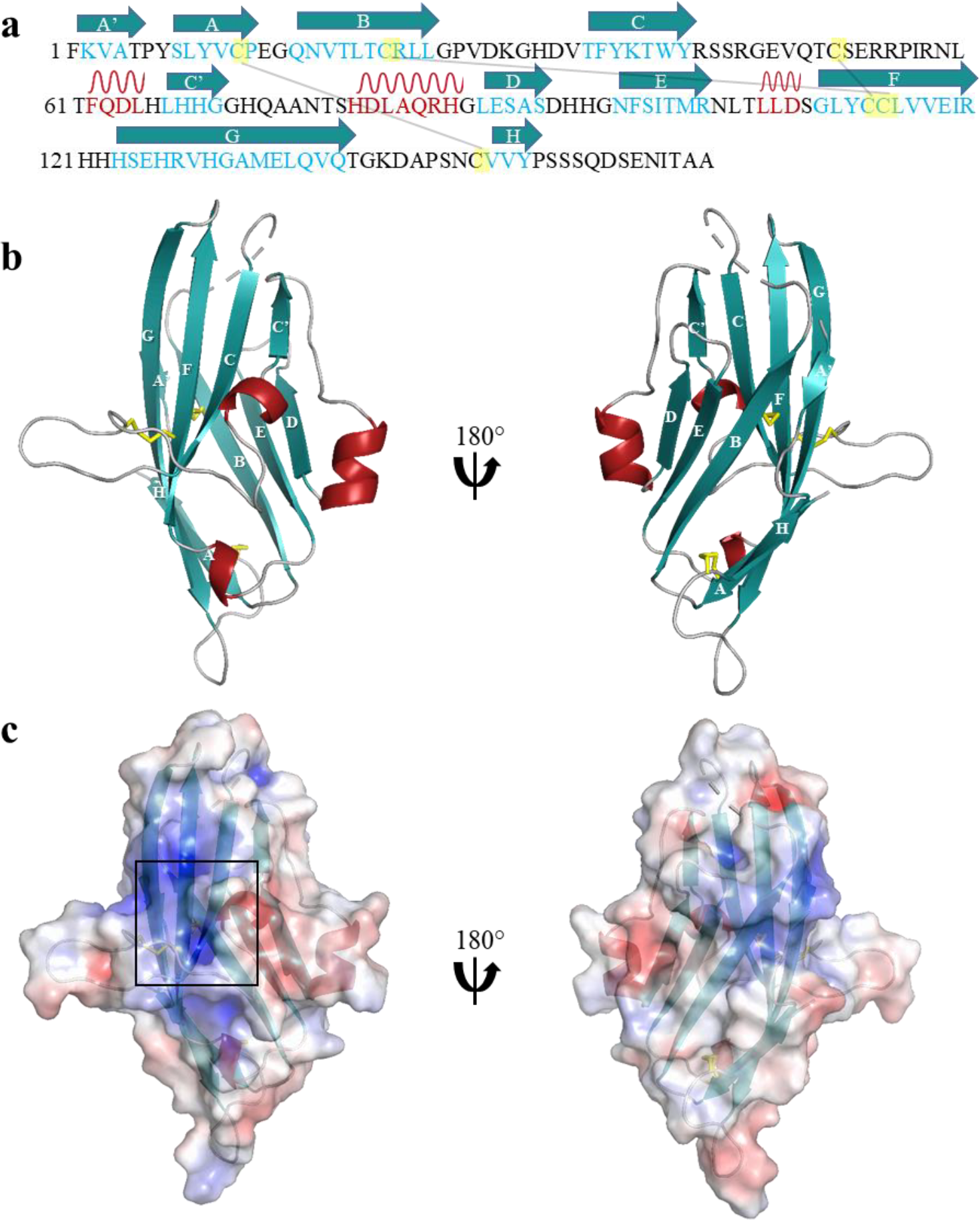
Human VISTA structure. **(a)** Sequence of human VISTA ECD with secondary structure and cysteines forming disulfide bonds indicated. **(b)** Cartoon structure of human VISTA ECD colored by secondary structure with beta strands labeled by Ig-domain nomenclature. **(c)** Electrostatic map of human VISTA (red = negative, blue = positive) revealing positively charged face of protein (black square).

### Comparison of VISTA with other B7 family proteins

The canonical fold of the B7 family is comprised of two distinct domains, an IgV domain with nine beta strands and an IgC domain with seven beta strands^16^. Typically, the IgC domain is proximal to the membrane while the IgV domain is distal and interacts directly with its cognate receptor. Of the seven B7 family proteins that have been crystallized, VISTA is the only family member that lacks an IgC domain. To highlight the structural distinctions of VISTA, we aligned the VISTA ECD with the IgV domain of human PD-L1 (PDB: 4Z18), its closest homolog in the B7 family (22% sequence identity). The overall beta sandwich fold is evident in both proteins with seven beta strands in VISTA aligning to corresponding strands in the PD-L1 structure (Fig. 2A). There are however, four key differences between VISTA and the classic B7 family fold. First, VISTA contains ten beta strands, instead of the nine that typically make up an IgV fold. Second, VISTA contains an extra helix (sequence FQDL) in place of a longer beta strand C’ (Fig. 2B). This helix is located in the predicted positively charged patch and may constitute a unique epitope that distinguishes VISTA binding interactions from its B7 homologs. Third, VISTA contains a 21-residue unstructured loop (C-C’ loop) that does not align with any B7 family structure (Fig. 2C). This region contains seven charged, surface exposed residues. PD-L1 and other B7 family proteins have a significantly smaller four residue loop at this location that directly connects two beta strands but does not protrude from the classic beta sandwich fold. Finally, VISTA also contains two additional disulfide bonds that are not present in any other B7 family protein (Fig 2D). These two disulfide bonds connect residues C12/C146 and C51/C113, respectively, while the conserved disulfide bond connects beta strands B and F (C22/C114). In particular, the C51/C113 disulfide bond connects the unique unstructured loop to the internal beta sandwich and likely plays a role minimizing flexibility in this region. The uniqueness of the VISTA structure was further corroborated by structural similarity comparisons with other B7 family proteins. The DALI server was used to calculate z-score similarities based on pairwise structural comparisons of known B7 family proteins (Fig. 2E). VISTA is most similar to PD-L1 (11.2) and PD-L2 (10.2), but displays larger structural differences with CD276 (7.1), CD80 (9.4), and CD86 (9.5), exemplifying its structural individuality among the B7 family. The average pair-wise z-score (mean of row in Fig. 2E) of each protein was compared against all other B7 family members. VISTA has an average pairwise z-score below 10 while other B7 family member averages are all 12 or higher.

**Figure 2.**
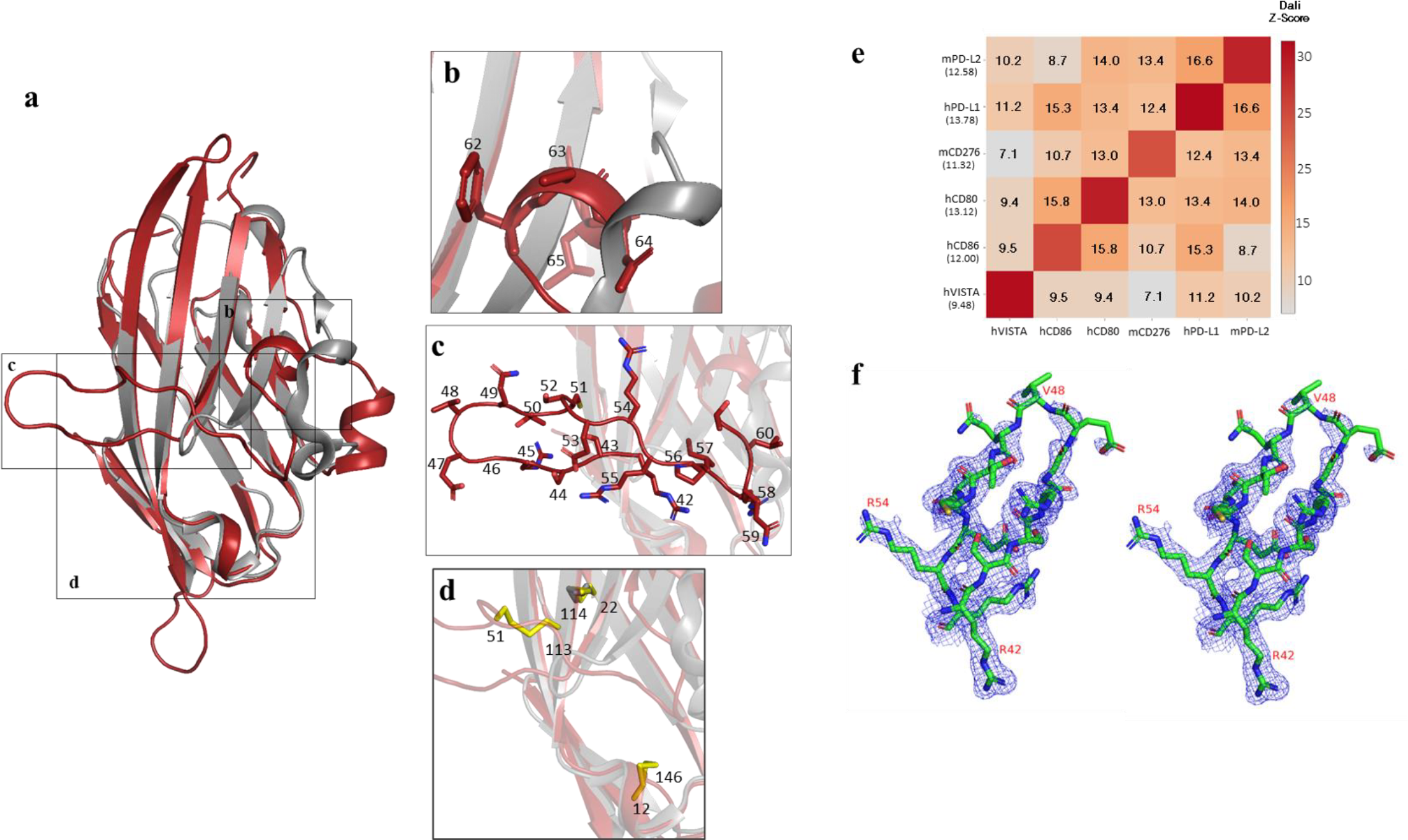
B7 family comparison. **(a)** Cartoon structure of human VISTA ECD (red) aligned with IgV domain of human PD-L1 (gray). **(b)** Unique helix in VISTA in place of beta strands in PD-L1. **(c)** Unique C-C’ loop in VISTA that extends from the beta-sandwich core. **(d)** Disulfide bonds in VISTA, including two unique disulfides in addition to conserved disulfide bond (C22, C114) between strands B and F. **(e)** Heatmap of Dali pairwise Z-scores between hVISTA and five other B7 family proteins. **(f)** Omit map of C-C’ loop.

The extended C-C’ loop was further examined for structural validity and uniqueness. An omit map was generated to verify loop density (Fig. 2F). Additionally, a DALI search of the C-C’ loop region uncovered homology with a protein known as immune receptor expressed on myeloid cells-1 (IREM-1). Analogous to VISTA, IREM-1 has an extended C-C’ loop held in place by a disulfide bond and also functions as a single domain inhibitory receptor on the surface of myeloid cells^17^. Even though the C-C’ loops differ in size and IREM-1 is not in the same predicted protein family as VISTA, a DALI z-score of 12.4 was calculated between VISTA and IREM-1, higher than any pairwise comparison of VISTA and B7 family proteins.

### Comparison of VISTA among species

When developing drug candidates, researchers often verify functionality by examining therapeutic efficacy and mechanism of action in preclinical murine tumor models, followed by toxicity studies in rodents and cynomolgus monkeys. Generating a species cross-reactive drug that binds its targets across multiple animal models allows for more facile transition between stages of development. Here, we compare the amino acid sequences and structural models of human, mouse, and cyno VISTA to highlight features helpful for directing development of cross-reactive drugs.

Using the crystal structure of human VISTA as a template, homology models of mouse and cyno VISTA were built using Rosetta. Three and nine-mer fragments of mouse and cyno VISTA were generated through the Robetta server, protein alignments were generated using Clustal Omega, and the Rosetta hybridize protocol was used to generate 10,000 potential structures of each target. These decoy structures were clustered and the lowest free energy structure from the largest cluster was used for structural comparison. Mouse and cyno VISTA homology models were aligned with the human VISTA crystal structure using PyMOL (Fig. 3A). Due to high sequence identity of human to mouse (70.4%) and human to cyno (96.4%), and the fact that human VISTA was used as a singular template for Rosetta hybridization, it is unsurprising that the proposed structures align very well to each other (RMSD of 0.592 and 0.430, respectively).

**Figure 3.**
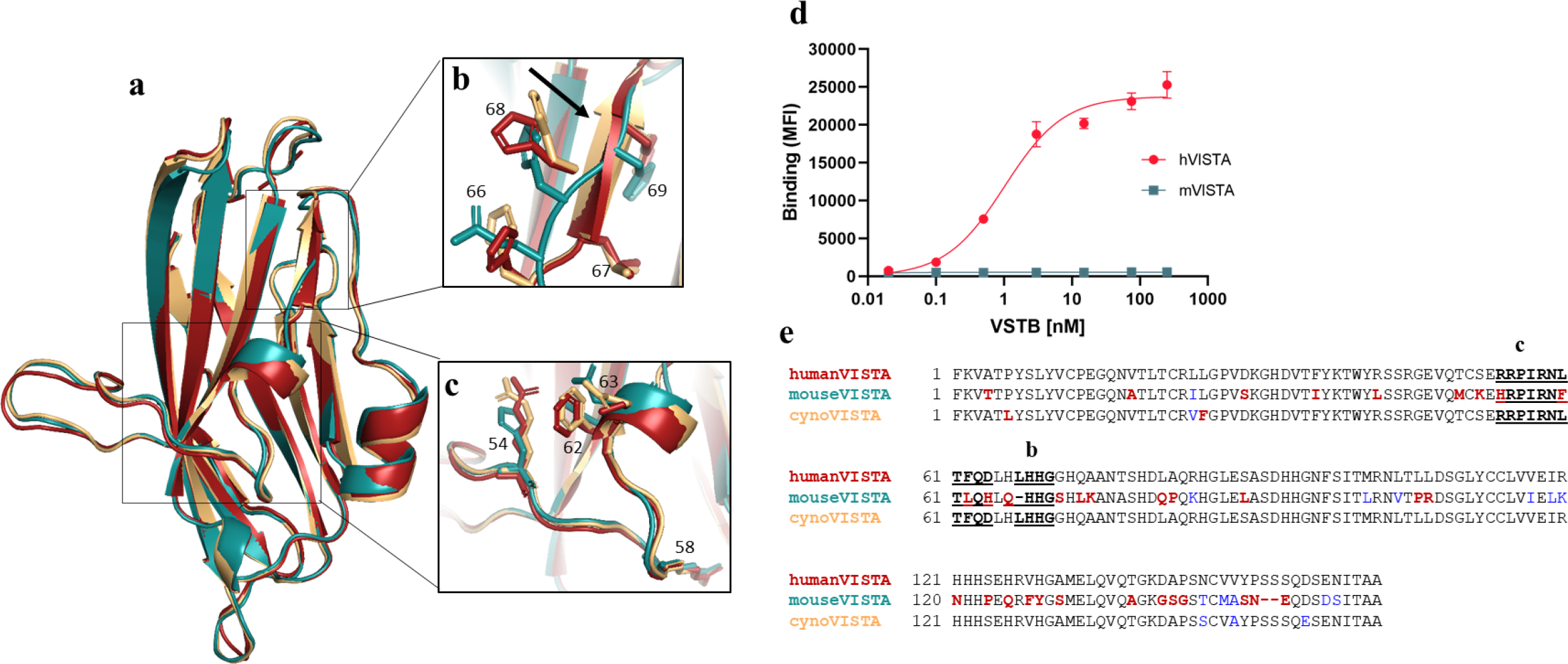
Comparison of human, murine, and cyno VISTA. (**a)** PyMOL Alignment of human VISTA ECD (red) and homology models of mouse VISTA (cyan) and cyno VISTA (beige). **(b)** Beta strand that is absent in mouse VISTA. Side chains shown as sticks, highlighting the lack of leucine at residue 66 and the deletion at residue 67 in the mouse VISTA sequence. **(c)** Portion of the C-C’ loop that is not conserved among species of VISTA. Residue differences in the mouse sequence are evident at positions 54 and 62. **(d)** Binding of VSTB antibody to human or mouse VISTA displayed on yeast. **(e)** Protein sequence alignment of human, mouse, and cyno VISTA. Residue differences from human VISTA (bold red), differences within same amino acid category (blue), unique beta strand (bold underlined, b) and unique loop+helix (bold underlined, c).

There are, however, several important structural distinctions between VISTA from different species. First, beta strand C’ (sequence LHHG) is only present in human and cyno VISTA (Fig. 3B). Mouse VISTA contains a H66Q mutation and a deletion of L67 that prevents formation of a beta strand in this position. As a result, mouse VISTA contains a total of 9 beta strands compared to the 10 found in human and cyno VISTA. Perhaps more importantly, the unique unstructured loop and helix region that comprise a positively charged face is modified in mouse VISTA (Fig. 3C). Within this charged region, mouse VISTA has four residues that differ from human VISTA: R54H, L60F, F62L, and D64H. The lack of two charged residues (R54 and D64) and the presence of two extra aromatic rings (H54 and H64) at these positions could play a role in altering charged electrostatic or pi-stacking interactions in this region. To confirm the importance of these structural differences for the molecular recognition of the VSTB antibody, equilibrium binding constants (K_d_) to mouse or human VISTA were measured (Fig. 3D). The K_d_ of the hVISTA/VSTB interaction is ~250 pM while the K_d_ of mVISTA/VSTB is undetectable due to lack of significant binding signal. The high degree of similarity between human and cyno VISTA and the increased divergence of mouse VISTA is exemplified in the multiple sequence alignment (Fig. 3E, bold/red). Highlighted differences between mouse and human VISTA in the unstructured loop and helix region likely underlie the lack of species cross-reactivity for the VSTB antibody.

### Mapping the VSTB/VISTA binding epitope

Next, we determined a putative binding epitope of an anti-VISTA antibody VSTB, which is a known inhibitor of VISTA signaling and prevents tumor growth in an MB49 mouse model of bladder cancer^15^. Fine-epitope mapping of VSTB was performed by screening a large library of VISTA mutants displayed on the surface of individual yeast cells to isolate variants that exhibited loss of antibody binding. Using this method^18^, we elucidated a set of VISTA residues that mediate VSTB binding.

A library of VISTA mutants was created via error prone-PCR using a low mutagenic rate to achieve, on average, a single amino acid mutation per gene. Restricting the library to single amino acid mutations allows for confident attribution of binding changes to a particular residue. A library with estimated diversity of 3.6×10^8^ yeast transformants was generated in a strain of *S. cerevisiae* engineered for surface protein display^19,20^. In addition to the yeast-displayed VISTA library and VSTB antibody, fine-epitope mapping of the VISTA/VSTB interaction required a control antibody to validate proper folding of VISTA mutants. The control antibody (referred to as ‘Ctrl’) was tested for conformational and distinct epitope binding. Heat denaturation of yeast-displayed VISTA followed by incubation with Ctrl antibody showed a lack of binding, confirming a conformational epitope that depends on VISTA structural integrity (Supp. Fig. 2a). Additionally, the Ctrl and VSTB antibodies were found to have distinct epitopes through the detection of simultaneous binding of both antibodies (Supp. Fig. 2b) Screening for yeast-displayed VISTA mutants that decreased binding to VSTB but still bound the Ctrl antibody allowed the isolation of residues that directly altered VSTB binding without disrupting the structural integrity of VISTA.

The library was induced for VISTA expression on the cell surface, resulting in each yeast displaying thousands of copies of an individual VISTA variant. Iterative rounds of fluorescence-activated cell sorting (FACS) were used to select yeast-displaying VISTA mutants that either lost binding to VSTB (“negative” sort) or retained binding to the Ctrl antibody (“positive” sort) (Fig. 4A). Following each sort round, collected yeast were cultured, and cell surface display of VISTA was again induced prior to the next round of FACS. In Sort 1, the library was incubated with 10 nM VSTB and screened to isolate VISTA mutants that displayed moderate to negligible binding to VSTB. To increase stringency, yeast collected from Sort 1 were subject to a higher concentration of VSTB in Sort 2 to isolate VISTA mutants demonstrating even weaker binding to VSTB. A VSTB-negative binding population was clearly enriched in Sort 2 (25.2% in gate) compared to the small number of negative clones observed in Sort 1 (3.4% in gate). In Sort 3, 50 nM of Ctrl antibody (about 200x the estimated K_d_) was used to isolate yeast-displayed VISTA mutants that retained structural integrity to bind Ctrl antibody. The screening stringency was again increased in Sort 4 by using an even higher concentration of VSTB (200 nM) to select for mutations that almost completely decreased antibody binding. By Sort 4, close to 70% of the yeast-displayed VISTA clones showed weak to negative binding to VSTB. Remaining yeast clones were subject to a final positive sort using a lower concentration of Ctrl (about 20x the estimated K_d_) to further confirm retention of VISTA structural integrity.

**Figure 4.**
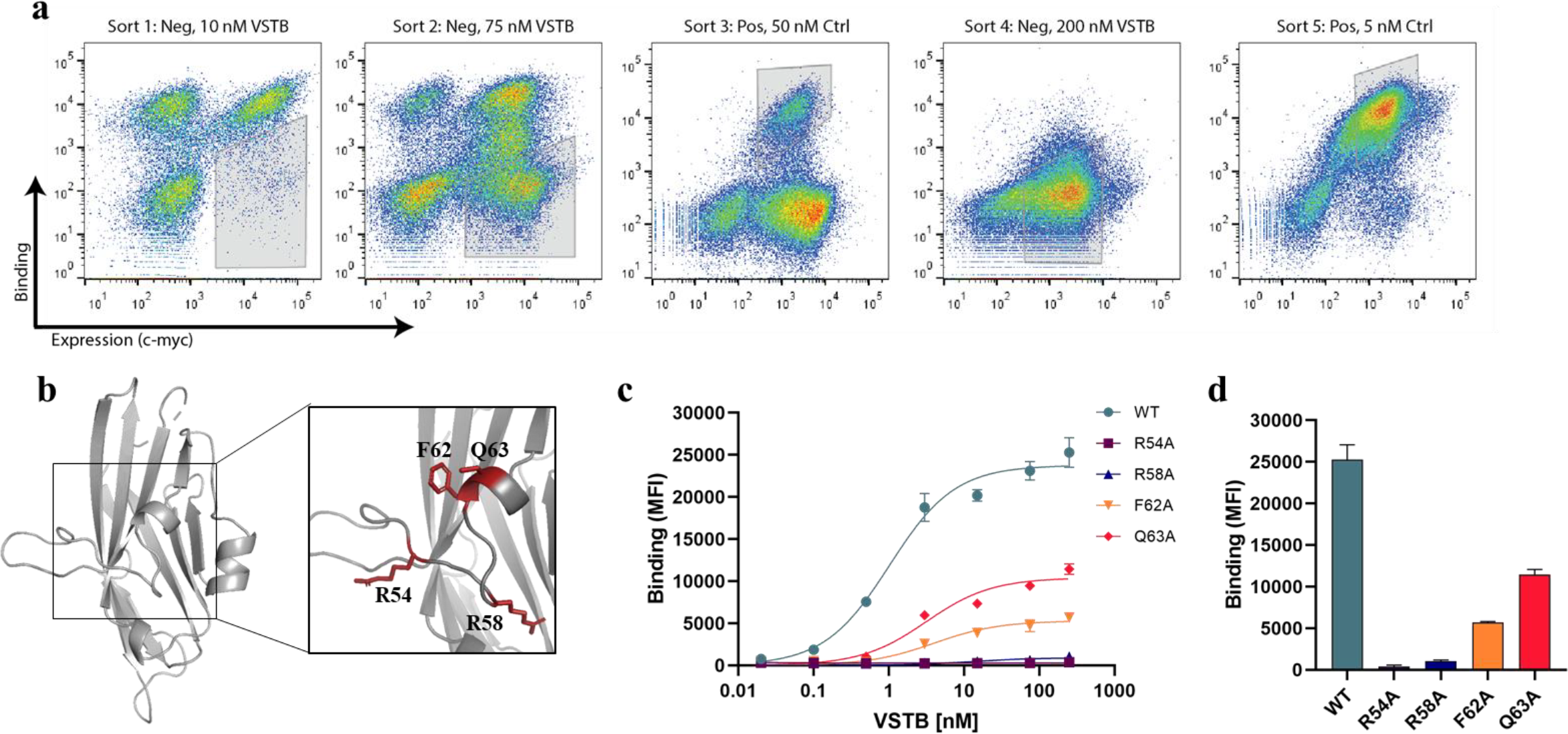
Mapping the VSTB binding epitope of VISTA. **(a)** Library screening used to isolate yeast-displaying VISTA mutants that retain or reduce antibody binding. FACS gates are shown in gray boxes on dot plots of individual positive or negative sorts. **(b)** Four residues identified from epitope mapping highlighted in red on the human VISTA ECD structure. **(c)** Binding of soluble VSTB antibody to yeast-displayed human VISTA with single amino acid alanine substitutions. **(d)** Binding signal comparison of 250 nM VSTB to yeast-displayed hVISTA alanine mutants.

Following library screening, 50 yeast clones were randomly selected for sequencing analysis to help identify a subset of residues directly involved in VSTB binding. Five mutations appeared in multiple (>4) sequenced clones: F62L, R54C, S124P, Q63R, and R58G (Supp. Table 1). All identified residues, with the exception of S124P, were localized to the alpha helix and unstructured loop fragment that is unique to the VISTA structure (Fig. 4B). Each residue was then individually mutated to alanine via site-directed mutagenesis to confirm that the residue locations, as opposed to the specific amino acid mutations, were drivers of VSTB binding. VISTA mutants were individually displayed on the surface of yeast and binding to VSTB was measured by flow cytometry. The binding of S124A to VSTB appeared to be equivalent to that of wild-type VISTA binding to VSTB (Supp. Fig. 3), suggesting that the proline mutation in S124P indirectly affected VSTB binding by modifying the local fold of the VISTA protein. This mutant was therefore excluded from the proposed epitope and was not subject to further analysis. In contrast, the remaining four mutants (R54A, R58A, F62A, or Q63A) were each found to individually affect binding to VSTB (Fig. 4C). Binding signal generated from 250 nM VSTB clearly shows weak binding of F62A and Q63A mutants and almost no detectable binding of R54A and R58A mutants, demonstrating that these four residues are critical for VISTA binding to VSTB (Fig. 4D).

### Mapping the VSIG3/VISTA interaction

The VSTB antibody has been shown to inhibit VISTA signaling, thus the residues we identified above suggest a functional epitope through which to guide future drug discovery efforts. Since the epitope for the recently-proposed VISTA ligand VSIG3 is unknown, we tested whether VSTB operates through direct ligand competition of VSIG3 at the residues mapped above and demonstrate overlap of the purported VSTB binding epitope with the VISTA/VSIG3 interaction.

Wild-type (WT) VISTA and a VISTA triple mutant containing the R54A, F62A, and Q63A mutations (Fig. 5A) were solubly expressed in mammalian cells. The R58 residue was not mutated in this analysis because structural examination revealed its importance to the stability of the entire loop due to its interior direction and proximity to other side chains. To confirm structural integrity of the triple mutant, the Ctrl antibody was first tested for binding to WT VISTA or the 54A/62A/63A triple mutant using an ELISA-based assay. Binding to the Ctrl Ab was retained for the 54A/62A/63A triple mutant with no significant difference in binding from WT VISTA in any concentration tested (Fig. 5B). In contrast, the three mutations significantly diminish binding to VSTB at every concentration tested (>95% decrease compared to WT VISTA). The R54A, F62A, and Q63A mutations therefore abrogate binding to VSTB but do not alter the VISTA structure significantly to abolish binding to the Ctrl antibody. A binding assay was then performed between VSIG3 ligand and the triple mutant or WT VISTA (Fig. 5C). The triple mutant showed a significant decrease in affinity for VSIG3, indicating that VISTA binding to VSIG3 is highly dependent on three of the same mutations that comprise the VSTB binding epitope. To further confirm the shared epitope, we pre-complexed WT VISTA with varying concentrations of VSTB and measured binding to VSIG3 (Fig. 5D). Dose-response disruption of VSIG3 binding is evident, with concentrations above 500 nM VSTB completely abolishing the VISTA/VSIG3 interaction. The VSIG3 binding signal of the 54A/62A/63A mutant and pre-complexed VISTA/VSTB is significantly lower than WT VISTA and pre-complexed VISTA/isotype control (Fig. 5E). This analysis suggests that the mapped VISTA epitope is not only important for interaction with an antibody that has been shown to inhibit VISTA signaling, but also drives binding to VSIG3, a known functional partner of VISTA.

**Figure 5.**
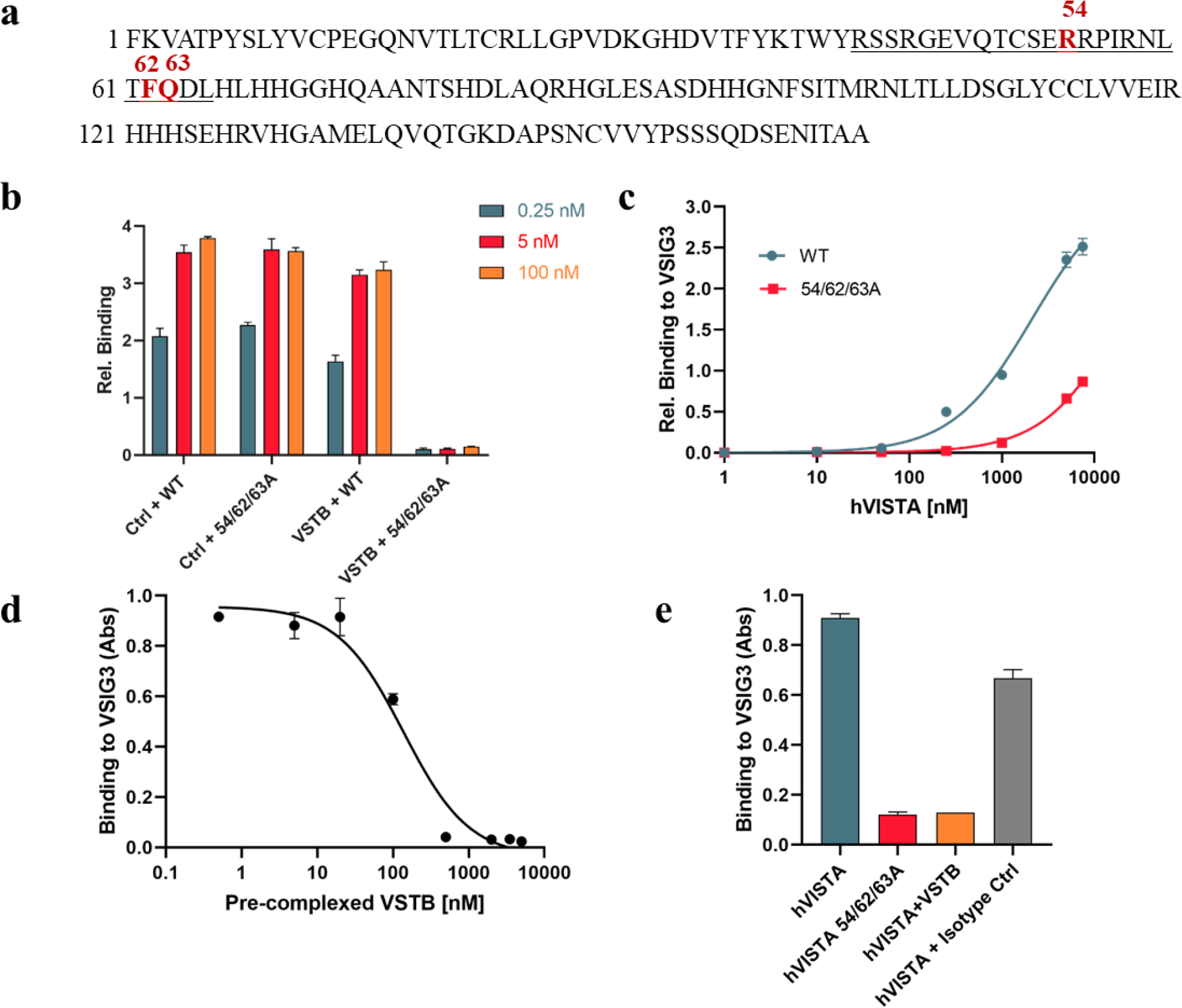
VSIG3 interaction with VISTA. **(a)** Sequence of hVISTA ECD with C-C’ loop underlined and mutated residues (54, 62, and 63) in red. **(b)** Binding of WT or VISTA 54A/62A/63A to Ctrl or VSTB antibody in ELISA format. VISTA was added to well coated antibody at three different concentrations. **(c)** Binding assay via indirect ELISA with well coated VSIG3 incubated with soluble WT or VISTA 52A/62A/63A. **(d)** Binding of 1 µM WT VISTA pre-complexed with a titration of VSTB concentrations to well coated VSIG3 in ELISA format. **(e)** VSIG3 binding signal comparison of 1 µM WT VISTA, 1 µM VISTA 54A/62A/63A, 1 µM VISTA pre-complexed with 500 nM VSTB, and 1 µM VISTA pre-complexed with an IgG isotype control.

## Discussion

In this work, we determined the structure of VISTA at a high resolution using multiple protein deglycosylation strategies and a combinatorial MR-Rosetta pipeline to solve the final structure. A combination of genetic mutations (N->Q) in the VISTA sequence, a glycosylation inhibitor added to the mammalian cell culture media, and enzymatic deglycoylsation post-purification was required to produce VISTA at its expected molecular mass. Previous crystallization attempts from our group using wild-type VISTA or Endo Hf-treated VISTA without genetic mutations were not successful, highlighting the importance of these steps for crystal growth. Additionally, solving the structure required the structural modeling efforts via Rosetta and an iterative pipeline of structure refinement and molecular replacement. In particular, the loop and helix regions that do not fit the classic Ig-like fold could not be solved with molecular replacement alone and required a Rosetta-based design approach. We foresee the strategy described here and the deposited VISTA coordinates assisting future efforts in solving difficult Ig-like protein structures.

A combinatorial strategy for fine-epitope mapping was used to isolate a shared epitope important for binding to VSTB, an anti-VISTA antibody of therapeutic interest, and this information in turn was used to determine a proposed overlapping epitope for the VSIG3 ligand. Previously, hydrogen-deuterium exchange was used to highlight a number of potential binding hotpots of VSTB to VISTA^15^. In our work, we provide clarity on the most important region of interaction and provide evidence corroborating the relevance of this epitope for VISTA function. We recognize that the yeast display-based epitope mapping approach used here may not elucidate every region involved in binding. However, because these mutations appeared with the highest frequency in our combinatorial library screen and the fact that each isolated residue mutated to alanine disrupts binding, the likelihood is high that this is the critical binding epitope that drives VISTA/VSTB interaction. VSIG3 was only recently discovered by ELISA-based binding screens as a cognate binding partner for VISTA expressed on T cells^5^. Here, we build on this evidence by confirming ELISA-based binding of VSIG3 to VISTA and show that the antagonist antibody VSTB blocks this specific interaction. Although there may be other VISTA binding partners and inhibitory antibodies may partially function through induced allosteric change of VISTA, we hypothesize that the inhibitory function of the VSTB antibody is at least partly due to its blocking of the VSIG3/VISTA binding, analogous to anti-PD-1 antibodies blocking the PD-1/PD-L1 signaling axis^21^. Future experiments using antibodies that bind VISTA but do not block VISTA/VSIG3 binding will be needed to confirm the importance of this signaling axis for anti-VISTA immunotherapy.

Comparing the structure of VISTA to PD-L1 reveals important qualities that diverge from the B7 protein family. Previously, sequence alignment of VISTA and PD-L1 highlighted a set of two conserved cysteines (C22/C114) that form the characteristic disulfide bond found in Ig-like proteins^1^. Examining the crystal structure of VISTA reveals two extra disulfide bonds that could not be confirmed with sequence analysis alone. Sequence analysis also isolated an unaligned region between strands C and C’ (Supp. Fig. 4), but structural data was needed to represent this as a solvent-exposed, flexible loop and helix. This extended C-C’ loop is unique among the B7 family, but shares homology with the inhibitory receptor IREM-1. The loop extends outwards from the beta-sandwich core and could play a role in promoting dimerization with another VISTA molecule, as in the case of growth factor receptor dimerization such as that observed with EGFR^22^. Alternatively, the loop could disrupt VISTA dimerization by preventing intermolecular interactions between Ig-like domains such as those found in PD-L1 dimers^23,24^. Further work via targeted deletion of the region and downstream functional analysis is needed to elucidate the role of the C-C’ loop in VISTA signaling. Additionally, VISTA contains a singular IgV-like domain while all other B7 family members contain both an IgV and an IgC domain. The B7 family members B7-1 (CD80), B7-2 (CD86), B7-DC (PD-L2), B7-H1 (PD-L1), and B7-H3 (CD276) are all dual-domain proteins and all function primarily as ligands. In contrast, the cognate receptors of these proteins including CD28, CTLA-4, ICOS, and PD-1 all have single IgV domain structures. Based on domain composition, VISTA appears to be more similar in architecture with the receptors rather than the B7 family ligands. Even though VISTA has shown functionality as a ligand in T cell proliferation assays^2^ and as a soluble Fc-fusion drug for autoimmune disease^25^, its structural composition as a single IgV domain and its binding interaction with VSIG3 point to its functionality as a receptor. Corroborating its functionality as a receptor on T cells, VISTA knockout T cells (Vsir ^-/-^) have been shown to proliferate more than wild-type cells in response to antigen-presenting cells *in vitro*^3^, and an agonistic anti-VISTA antibody (MH5A) reduced the allogeneic T cell response in a murine model of graft-versus-host disease^4^.

Mouse and human VISTA share 70.4% sequence identity but have important structural differences. Through sequence analysis alone, the two proteins were predicted to have very similar folds due to conserved cysteines as well as a lack of significant gaps in the alignment^1^. Structural comparisons between human VISTA and a Rosetta-based homology model of mouse VISTA reveal critical structural differences in the fragments surrounding the VISTA epitope. We propose that the lack of a beta strand at residues 67-70 and the differences in the epitope helix (FQDL→ LQHL) cause side chain orientation changes that directly prevent VSTB from being cross-reactive with murine VISTA. The differences in this critical region suggest that inhibitor drugs binding to the mapped epitope will be cross-reactive between human and cyno VISTA, which exhibit a high degree of similarity, but will likely not bind to mouse VISTA.

Knowledge of the three-dimensional structure of VISTA and residues that comprise its binding epitope can help guide future drug development by enabling small molecule library screening through computer-aided drug design (CADD)^26^ and computational antibody screens through antibody-antigen docking^27,28^. Additionally, the high resolution structure can support future studies of receptor or ligand interactions through computational docking experiments. The coordinates for the VISTA ECD provided here will also expedite co-crystallization efforts of VISTA complexes by providing a well-suited template for molecular replacement. The initial success of checkpoint inhibitors in the clinic has provided a blueprint for new drugs that release the breaks on the immune system. VISTA inhibitors have the potential to provide an orthogonal method of T cell stimulation and anti-tumor activity by directly affecting the APC/T cell signaling axis. We believe the high resolution crystal structure of VISTA presented here will bolster these efforts by encouraging further VISTA-related research and by directly assisting drug development endeavors.

## Materials and Methods

### Preparation of recombinant VISTA protein

The human VISTA extracellular domain (ECD) sequence with native signal peptide (Met1-Ala194, UniProt) was ordered as a gblock Gene Fragment (IDT) and cloned into the cytomegalovirus-driven adenoviral shuttle vector pAdd2 using standard Gibson cloning at EcoRI/XhoI vector cut sites. Protein was expressed in Expi293 cells according to the manufacturer’s protocol, and proteins were purified from culture supernatant using nickel affinity chromatography. A hVISTA triple mutant (R54A, F62A, Q63A) used for epitope binding verification was produced in a similar manner. For crystallization, an asparagine triple mutant (N59Q, N76Q, N158Q) was cloned into the pAdd2 expression plasmid as described above and expressed in Expi293 cells in the presence of 10 µM Kifunensine (Cayman Chemical, 109944-15-2). After nickel affinity chromatography, N-linked glycans were removed using endoglycosidase H (Endo Hf, New England BioLabs, P0703). De-glycosylated VISTA protein was separated from Endo Hf via additional nickel affinity chromatography. Residues are numbered starting after the signal peptide (Phe1, Lys2, Val3).

### Crystallization and data collection

VISTA ECD protein was concentrated to 8 mg/mL and buffer exchanged into 50 mM HEPES (pH 8.2), 50 mM NaCl for crystallization trials. Initial crystals were grown at 12° C by mixing the protein solution with equal volume of reservoir solution (0.2 M NaBr and 20% PEG 3350). The diffraction analysis showed poor multiple diffraction spots to around 4Å. Fine tuning attempts using various additives and detergents did not improve the crystal morphology. Since crystal morphology at 12 °C and 20 °C were similar, further optimization attempts were performed at 20 °C. A grid search using various buffers identified HAT (made by mixing equal volumes of 1M Tris (pH 8.0), 1M HEPES (pH 7.5), and 1M ADA (pH 6.5)) as the optimal buffer for crystal formation. Seeding protocols with 1:1000 diluted crystal seeds introduced to the drop after two days gave small single crystals. A grid search by varying concentration of PEG and NaBr and also varying the drop ratio generated the best crystals with a well solution containing 75 mM NaBr, 18% PEG 3350, and 50 mM HAT buffer. The drop ratio for the best crystals was 0.8 µL of protein and 0.6 µL of well solution.

The crystals were flash cooled by dipping in a well solution containing 32% PEG 3350. Diffraction data sets were collected at 100° K via Stanford Synchrotron Radiation Lightsource (SSRL) beamline 12-2 at a wavelength of 0.98 Å using PILATUS 6M detector. Data were indexed and integrated using the XDS package^29^. The crystals belong to space group P2_1_ and contain one monomer per asymmetric unit. The best crystals diffracted to around 1.7 Å and the final data (480 degrees) is processed to 1.85 Å resolution. In addition, low dose and highly redundant sulfur SAD data were collected at a wavelength of 1.55 Å by using the 5 degree inverse beam geometry (total 4670 degrees of data). The crystallographic data are summarized in Table 1.

**Table 1:** Data collection and refinement statistics

|  | VISTA |
| --- | --- |
| **Data collection** |  |
| Space group | P2<sub>1</sub> |
| Cell dimensions |  |
| *a*, *b*, *c* (Å) | 29.17, 64.09, 37.69 |
| $\alpha$, $\beta$, $\gamma$ (°) | 90.0, 91.7, 90.0 |
| Resolution (Å) | 40-1.85 (1.90-1.85) * |
| *R*<sub>sym</sub> or *R*<sub>merge</sub> | 10.1 (156.2) |
| *I* / $\sigma I$ | 13.0 (1.5) |
| Completeness (%) | 98.4 (96.2) |
| Redundancy | 9.2 (8.4) |
| **Refinement** |  |
| Resolution (Å) | 40-1.85 |
| No. reflections | 11694 |
| *R*<sub>work</sub> / *R*<sub>free</sub> | 0.18/0.22 |
| No. atoms |  |
| Protein | 1219 |
| Ligand/ion | 0 |
| Water | 41 |
| *B*-factors (Å<sup>2</sup>) |  |
| Protein | 41.2 |
| Ligand/ion |  |
| Water | 43.3 |
| R.m.s. deviations |  |
| Bond lengths (Å) | 0.006 |
| Bond angles (°) | 1.002 |

\*Values in parentheses are for highest-resolution shell.

### Structure determination and refinement

The human VISTA ECD sequence was submitted to the GREMLIN server (gremlin.bakerlab.org) for co-evolution contact predictions and a list of top homology models. Ten thousand decoy structures of hVISTA were generated using Rosetta homology modelling (RosettaCM) with the top ten VISTA homologs obtained from the GREMLIN search as templates. The initial screen running Phaser^30^ for possible molecular replacement solutions was conducted on the 10,000 structures generated from the RosettaCM run, but failed to produce definitive hits. Model convergence of the top 100 scoring RosettaCM solutions was analyzed to produce a partial consensus model (88 amino acids). An MR solution was obtained using the partial model. The model generated by MR search and Phenix^31^ building resulted in a reliable structure with Rwork and Rfree of 0.48 and 0.54, respectively. Various building attempts with Phenix, Phenix-rosetta, RosettaRemodel (for generating templates for VISTA specific disulfides), and Buccaneer^32^ programs brought the Rwork and Rfree to 0.37 and 0.41, respectively. Then, Phenix autobuilding by incorporating the sulfur SAD data, Buccaneer building, and extensive manual building resulted in Rwork and Rfree of 0.26 and 0.31, respectively. Further refinements were done by using Phenix and manual model building. The final model includes one protein monomer, two NAG molecules, and 41 water molecules in the asymmetric unit. No electron density was observed for the residues 29 – 31, and for the C-terminal residues starting from residue 152. The refinement converged to a final Rwork and Rfree of 0.18 and 0.22, respectively. There are no Ramachandran outliers and 98.6% of the residues are in the favored region. The refinement statistics are provided in Table 1. All crystal structure figures were created using PyMOL.

### Homology modeling and structural comparisons

Structural models of murine VISTA ECD (Phe33-Ala191, Uniprot), cynomolgous monkey (cyno) VISTA ECD (NCBI# XP_005565644, Phe33-Ala194), and human VSIG3 ECD (Lys23-Gly241, Uniprot) were made using RosettaCM homology modeling, as described previously^33^. A single template of human VISTA was used for mouse and cyno VISTA modeling and an eight template set was used for VSIG3 modeling (PDB IDs: 1NBQ_B, 3JZ7_A, 3R4D_A, 4Z18_A, 5JHD_A, 5TEZ_I, 6CPH_D, 6EH4_D). 10,000 decoy structures of mouse VISTA, cyno VISTA and human VSIG3 were generated. The top 100 scores (lowest free energy) for each target were sorted using the standard Rosetta ‘cluster’ application and the top score of the largest cluster was used for downstream structural comparisons.

Structural alignments between VISTA species were performed using both the pymol ‘super’ and ‘align’ commands. Structural comparisons between the B7 family proteins and human VISTA were performed using the ‘all against all’ option in the online DALI server^34^ (http://ekhidna2.biocenter.helsinki.fi/dali/). The PDB IDs for the B7 family structures used in the comparison were CD80: 1DR9, CD86: 5YXK, PD-L2: 3BP5, PD-L1: 4Z18, CD276: 410K.

### Epitope mapping via library generation and screening

DNA encoding the human VISTA ECD amino acids (Phe33-Ala194, Uniprot), was cloned into the pCT yeast display plasmid^35,36^ using standard Gibson cloning. An error prone library was created using WT hVISTA as a template, and mutations were introduced using low-fidelity Taq polymerase (Invitrogen) and nucleotide analogs 8-oxo-dGTP and dPTP (TriLink Biotech) as described previously^37,38^. Three different PCR reactions of 15 cycles were performed with 1.25, 1.5, and 1.75 µM of dNTP analogs. The 1.75 µM library was found to have the highest percentage of single amino acid mutations. This library was amplified and purified using gel electrophoresis. Empty pCT vector was cut using NheI and BamHI restriction sites. The amplified insert and cut vector were electroporated in a 5:1 DNA weight ratio into EBY100 yeast, where they were assembled *in vivo* through homologous recombination. Library size was determined to be 3.6×10^8^ by dilution plating.

The VSTB antibody used for screening was derived from the Janssen VSTB174 sequence^15^. We paired the VSTB174 heavy chain variable domain with the hIgG1 constant domain (Ala1-Lys330, Uniprot P01857) and paired the VSTB174 light chain variable domain with the human kappa light chain constant domain (Arg1-Cys107, Uniprot P01834). Heavy chain and light chain were individually cloned into the pAdd2 expression vector using standard Gibson cloning. The positive control ‘Ctrl’ antibody variable domains were paired with murine IgG2a constant domains (HC: NCBI# AAA37906, kappa LC: NCBI# BAB33404). Both antibodies were expressed in Expi293 cells with a 1:1 weight ratio of heavy chain: light chain DNA using the manufacturer’s protocol. Antibodies were purified from the supernatant using protein A affinity chromatography.

Yeast displaying hVISTA mutants that lost binding to VSTB but retained binding to Ctrl were isolated from the library using fluorescence-activated cell sorting (FACS). Equilibrium sort rounds were performed in which yeast were incubated at 4 °C for 12 hr in phosphate-buffered saline (PBS) containing 1 mg/mL BSA with the following concentrations of antibody: Sort 1, 10 nM VSTB; Sort 2, 75 nM VSTB; Sort 3, 50 nM Ctrl; Sort 4, 200 nM VSTB; Sort 5, 5 nM Ctrl. After incubation with antibody, yeast were pelleted, washed, and resuspended in PBS+BSA with a 1:5000 dilution of chicken anti-c-Myc (Invitrogen, A21281) for 30 mins at 4 °C. Yeast were then washed and pelleted, and labeled on ice with 1:250 dilution of secondary antibodies for binding (anti-mouse 488, ThermoFisher A11059 or anti-human 647, ThermoFisher A21445) and expression (anti-chicken 647, abcam ab150171 or anti-chicken 488, ThermoFisher A11039). Labeled yeast were sorted by FACS using a BD Aria sorter (Stanford FACS Core Facility). Negative sort gates for sorts 1, 2, 4 and positive sort gates for sorts 3, 5 were drawn to isolate populations with desired binding characteristics. Following FACS sort 5, plasmid DNA was recovered from yeast using a Zymoprep kit (Zymo Research Corp), transformed into DH10b electrocompetent E. coli, and isolated using a GeneJET plasmid miniprep kit (ThermoFisher, K0503). Sequencing was performed by MCLAB (Molecular Cloning Laboratories).

### Binding assays

Single alanine mutants of human VISTA (R54A, R58A, F62A, Q63A, or S124A) were generated using site-directed mutagenesis according to a standard two-stage QuikChange PCR protocol^39^. PCR fragments were cloned into the pCT yeast surface display vector and individually transformed into EBY100 yeast. The genes for WT human VISTA (33-194, Uniprot) and mouse VISTA (33-191, Uniprot) were also cloned into pCT and transformed into yeast as described above. Binding assays were performed by mixing surface-displayed VISTA on yeast (~50,000 molecules/cell)^40^ with a titration of target antibody concentrations (VSTB or Ctrl) in individual eppendorf tubes. Binding reactions were incubated at 4 °C for 12 hr to allow interactions to reach equilibrium. Yeast were labeled with the same reagents using protocols as described for library sorts and analyzed by flow cytometry on a BD Accuri. Binding populations were gated using FlowJo software and geometric means of fluorescence were plotted against concentration and fit to a one-site specific binding curve on GraphPad Prism. Error bars represent standard deviation of the mean for duplicate measurements.

For enzyme-linked immunosorbent assays (ELISAs), recombinant proteins were immobilized on a 96-well flat bottom plate (Corning, CLS3595) by incubation at 4 °C for 12-16 hr. VSIG3 was coated at 15 µg/mL and VSTB and Ctrl antibodies were coated at 2 µg/mL in PBS. Wells were washed with PBS + 1% Tween-20 and then blocked with PBS + 2.5% milk powder + 2.5% BSA at room temperature for 2 hr. Soluble His-tagged VISTA protein (WT or Ala triple variant) was added at varying concentrations in PBS + 0.1% BSA + 0.1% Tween-20 and the plate was incubated at room temperature for 2 hr. For VSTB pre-complexed experiments, 1 µM of VISTA was incubated with 1 nM – 5000 nM of VSTB in individual eppendorf tubes overnight and then added to VSIG3-coated ELISA plates. Binding of VISTA was detected indirectly by first adding 1:750 rabbit anti-6-HIS (Bethyl, A190-114F) and then adding 1:7500 anti-rabbit-HRP (Novus Biologicals, NB7160). Substrate solution (1-Step Ultra TMB, ThermoFisher, 34028) was added, reaction was stopped after 15 min with 2M sulfuric acid, and absorbance at 450 nM was read on a microplate reader (Synergy H4, BioTek). Absorbance values of control wells with no coated protein were subtracted from sample wells and corrected values were plotted against VISTA concentration and fit to a one-site specific binding curve on GraphPad Prism. Error bars represent standard deviation of the mean for triplicate measurements.

## Acknowledgements

The authors thank Daniel Fernandez for crystallization assistance and advice, Ryan Kelly for providing materials and advice used in epitope mapping, Qingping Xu for providing the script used for automated MR search, Jenifer Brown and Lindsay Deis for initial crystallization attempts and assistance, Sean Hunter and James Kintzing for helpful discussions regarding manuscript preparation, Jun Kim for providing helpful protein de-glycosylation strategies, the Stanford FACS Core Facility for assistance with flow cytometric sorting, and the ChEM-H Macromolecular Structure Knowledge Center (MSKC) for assistance with crystal screening. Portions of this research were performed at the Stanford Synchrotron Radiation Laboratory, a national user facility operated by Stanford University on behalf of the U.S. Department of Energy, Office of Science, and Office of Basic Energy Sciences under Contract No. DE-AC02-76SF00515. The SSRL Structural Molecular Biology Program is supported by the DOE Office of Biological and Environmental Research, and by the National Institutes of Health, National Institute of General Medical Sciences (including P41GM103393). This work was supported by the Stanford Bio-X Seed Grant program (J.R.C.), NSF Graduate Research Fellowship (N.M.), Stanford Graduate Fellowship (N.M.), Stanford Bio-X Undergraduate Summer Research Program (S.M.), and Stanford NIST Predoctoral Training Grant Program (A.P.S.).

## Contributions

N.M and S.M performed the protein preparation and initial crystallization attempts. I.I.M performed the crystallization optimization and x-ray diffraction. P.S.H. and I.I.M performed the structure refinement and determination. A.P.S. carried out homology modelling. N.M and S.M completed epitope mapping and N.M performed single clone analysis and functional epitope experiments. N.M. analyzed the data. N.M. and J.R.C. prepared the manuscript with input from all coauthors.

