## Supplementary Information for "Structure and Functional Binding Epitope of V-domain Ig Suppressor of T-cell Activation (VISTA)"

### Supplementary Figure 1

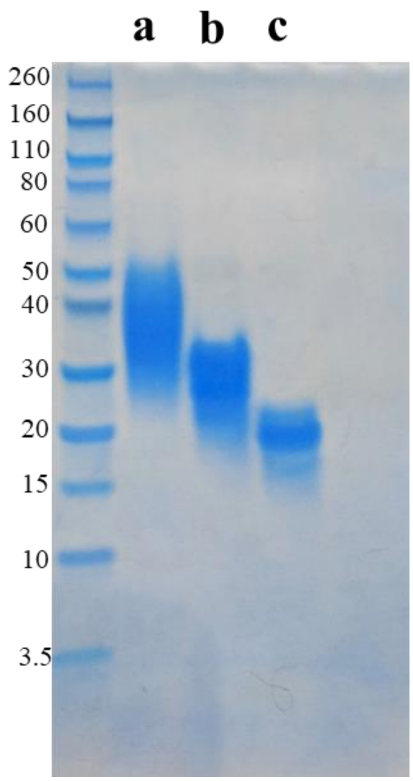


**Deglycosylation of VISTA.** SDS-PAGE gel of VISTA ECD at different stages of deglycosylation. **(a)** Wild-type VISTA (Met1-Ala194), **(b)** VISTA with three asparagine to glutamine mutations (N59Q, N76Q, N158Q), **(c)** VISTA with 3 N→Q mutations, Kifunensine in culture media, and Endo Hf enzymatic cleavage. The predicted molecular mass of VISTA ECD is 19 kDa. Only the combination of genetic mutations and enzymatic cleavage produced a distinct band at the estimated molecular mass.

**Supplementary Figure 2**

**a**

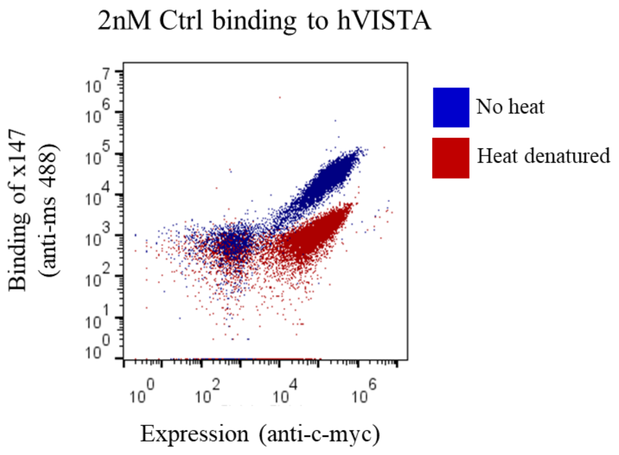


**b**

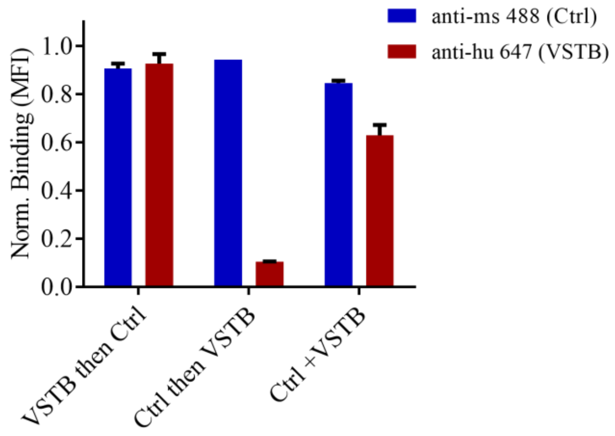


**Antibody verification for epitope mapping. (a)** Flow cytometry plot of Ctrl antibody binding to yeast-displayed hVISTA with and without heat denaturation of yeast. The decreased binding after heat denaturation confirms conformational (rather than linear) epitope binding. **(b)** Relative binding plot of Ctrl and VSTB antibodies to yeast-displayed hVISTA. Antibodies were added sequentially, where the first antibody was allowed to reach equilibrium, then the second antibody was added for 15 min, or together where both antibodies were allowed to reach equilibrium. Binding of Ctrl antibody = blue bar; binding of VSTB antibody = red bar, normalized to binding signal when antibodies are bound individually. The VSTB antibody does not preclude Ctrl antibody binding, while Ctrl antibody binding blocks the binding of VSTB. As shown in Ctrl+VSTB, it is possible for both antibodies to bind simultaneously, indicating distinct epitopes. Error bars represent standard deviation of the mean fluorescence intensity for triplicate measurements.

**Supplementary Figure 3**

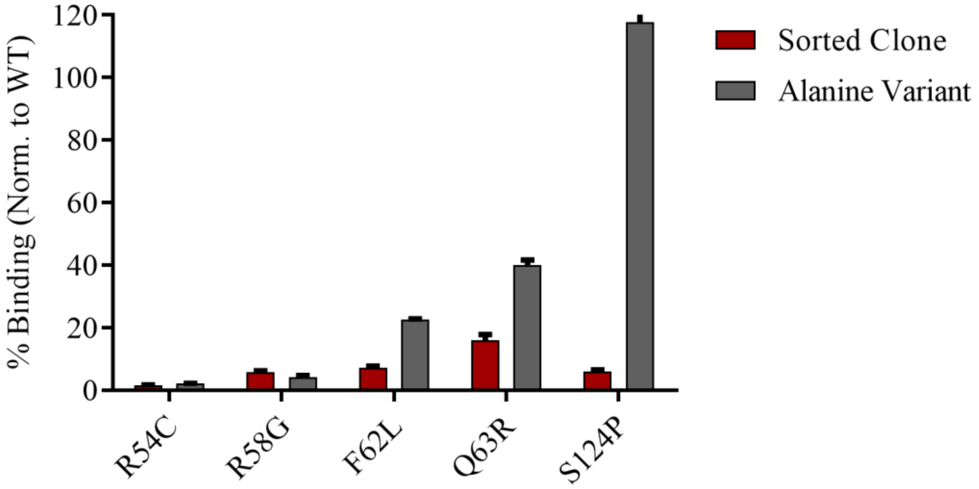


**Point mutant binding to VSTB.** The five hVISTA mutations identified from screening (red) and alanine variants at the same position (gray) were displayed as individual clones on yeast and measured for binding to VSTB (200 µM). Binding was normalized to wild-type (WT) hVISTA binding to 200 µM VSTB. S124A showed binding equivalent to WT VISTA levels, while all other hVISTA mutants and alanine variants showed a strong reduction in VSTB binding, suggesting these residues are involved in VSTB binding. Error bars represent standard deviation of the mean fluorescence intensity for triplicate measurements.

### Supplementary Figure 4

```
B7-H5(VISTA)     1 ----------FKVATPYSLYVCPEGQNVTLTCRLLGPVDKGHDVTFYKTWYRSSRGEVQT
B7-1(CD80)       1 ----------------VIHVTKEVKEVATLSCGHNV-SV-EELAQTRIYWQKEKKMVLTM
B7-2(CD86)       1 ---------------APLKIQAYFNETADLPCQFANSQN-QSLSELVVFWQDQENLVLNE
B7-DC(PD-L2)     1 ---------LFTVTVPKELYIIEHGSNVTLECNFDTGSH-VNLGAITASLQ---------
B7-H1(PD-L1)     1 ----------FTVTVPKDLYVVEYGSNMTIECKFPVEKQ-LDLAALIVYWEMEDKNIIQF
B7-H2(ICOSL)     1 -------------DTQEKEVRAMVGSDVELSCACPEGSR-FDLNDVYVYWQTSESKTVVT
B7-H3(CD276)     1 ---------AVEVQVPEDPVVALVGTDATLRCSFSPEPG-FSLAQLNLIWQLTDTKQ--L
B7-H4(VTCN1)     1 LIIGFGISGRHSITVTTVASAGNIGEDGILSCTFEP--D-IKLSDIVIQWLKEGVLGLVH

B7-H5(VISTA)    51 CSERRPIRNLTFQDLHLHHGGHQAANTSHDLAQRHGLESAS--DHHGNFSITMRNLTLLD
B7-1(CD80)      43 MS--------------------GDMNIWPEYKNRTIFD------ITNNLSIVILALRPSD
B7-2(CD86)      45 VY----------------LGKEKFDSVHSKYMGRTSFDSD-------SWTLRLHNLQIKD
B7-DC(PD-L2)    42 ----------------------KVENDTSPHRERATLLEEQ--LPLGKASFHIPQVQVRD
B7-H1(PD-L1     50 VH-----------------GEEDLKVQHSSYRQRARLLKDQ--LSLGNAALQITDVKLQD
B7-H2(ICOSL)    47 YH---------------IPQNSSLENVDSRYRNRALMSPAG--MLRGDFSLRLFNVTPQD
B7-H3(CD276)    49 VH-----------------SFTEGRDQGSAYANRTALFPDL--LAQGNASLRLQRVRVAD
B7-H4(VTCN1)    58 EF---------------KEGKDELSEQDEMFRGRTAVFADQ--VIVGNASLRLKNVQLTD

B7-H5(VISTA)   109 SGLYCCLVVEIRHHHSEHRVHGAMELQVQTGKDAPSNCVVYP--SSSQDSENITAA
B7-1(CD80)      77 EGTYECVVLKYEKDAFKREHLAEVTLSVKADFPTPSISDF---EIPTSNIRRIICS
B7-2(CD86)      82 KGLYQCIIHHKKPTGMIRIHQMNSELSVLANFSQPEIVPI--SNITENVYINLTCS
B7-DC(PD-L2)    78 EGQYQCIIIYGVAWDYK-----YLTLKVKASYRKINTHILKV---PETDEVELTCQ
B7-H1(PD-L1)    91 AGVYRCMISYGGA-DYK-----RITVKVNAPYNKINQRILVVD--PVTSEHELTCQ
B7-H2(ICOSL)    90 EQKFHCLVLSQSL-GFQEVLSVEVTLHVAANFSVPVV---SAPHSPSQDELTFTCT
B7-H3(CD276)    90 EGSFTCFVSIRDF-GSA-----AVSLQVAAPYSKPSMTLEPNKDLRPGDTVTITCS
B7-H4(VTCN1)   101 AGTYKCYIITSKGKGNA-----NLEYK-TGAFSMPEVNVD-----YNASSETLRCE
```

**B7 Family Sequence Alignment.** Multiple sequence alignment of human B7 family extracellular domains generated from Clustal Omega (https://www.ebi.ac.uk/Tools/msa/clustalo/). Sequences are truncated after the end of the human VISTA ECD sequence. Amino acid matches to VISTA are shown in red and residue similarities to VISTA (same subtype of side chain) are shown in blue.

### Supplementary Table 1

| Mutation | # of Samples |
|---|---|
| F62L | 10 |
| R54C | 9 |
| S124P | 9 |
| Q63R | 5 |
| R58G | 4 |
| L67P | 3 |
| F1L | 2 |
| S8P | 2 |
| C51R | 2 |
| F62S | 2 |
| D64G | 2 |
| E118G | 2 |
| C113R | 2 |

**Epitope mapping sequencing results.** Table of repeat mutations found from sequencing DNA isolated from the yeast population after Sort 5 of epitope mapping. The top five mutations by frequency are highlighted in yellow. Mutations that appeared four times or more were selected for single clone analysis and site-directed mutagenesis.
